# Modeling of DNA replication in rapidly growing bacteria with one and two replication origins

**DOI:** 10.1101/354654

**Authors:** Renata Retkute, Michelle Hawkins, Christian J. Rudolph, Conrad A. Nieduszynski

## Abstract

In rapidly growing bacteria initiation of DNA replication occurs at intervals shorter than the time required for completing genome duplication, leading to overlapping rounds of replication. We propose a mathematical model of DNA replication defined by the periodicity of replication initiation. Our model predicts that a steeper gradient of the replication profile is to be expected in origin proximal regions due to the overlapping rounds of synthesis. By comparing our model with experimental data from a strain with an additional replication origin, we predict defined alterations to replication parameters: (i) a reduced fork velocity when there were twice as many forks as normal; (ii) a slower fork speed if forks move in a direction opposite to normal, in line with head-on replication-transcription collisions being a major obstacle for fork progression; (iii) slower cell doubling for a double origin strain compared to wild-type cells; and (iv) potentially an earlier initiation of replication at the ectopic origin than at the natural origin, which, however, does not a˙ect the overall time required to complete synthesis.

## Introduction

Eukaryotic DNA replication is highly regulated to ensure that replication origins are activated only once during each synthesis phase (Blow and Gillespie, 2008). In contrast, bacteria can, under favorable conditions, increase their growth rate by permitting overlapping rounds of replication (Nordstrom and Dasgupta, 2006). Concurrent rounds of replication have also been shown to occur in some archaea with multiple replication origins (Hawkins *et al*, 2013). The majority of bacteria, including *Escherichia coli*, have one circular chromosome and a single replication origin, termed *oriC* (Duggin *et al*, 2008; Rocha, 2008). Two replication forks are recruited at *oriC* and proceed bidirectionally until they meet in an area opposite the origin. This area is flanked by multiple *ter* sites, which, if the *Tus* terminator protein is bound, forms a replication fork trap that allows forks to enter but not to leave (Rocha, 2008). This divides the *E. coli* chromosome into two defined halves or replichores. Progression of forks within each replichore is not necessarily smooth. Replication forks may be blocked or displaced from the chromosome when they encounter obstacles such as DNA lesions, proteins bound to DNA or transcribing RNA polymerase complexes. The majority of highly transcribed genes are oriented co-directionally with replication fork movement, indicating that head-on collisions of replication and transcription complexes might be particularly problematic (Brewer, 1988; McLean *et al*, 1998; McGlynn *et al*, 2012).

Cells undergoing rapid growth may initiate multiple, concurrent rounds of DNA replication such that each cell may contain two or more origins at the time when septation is complete (Fig 1A). This allows cells to grow with a doubling time less than that required to copy the whole genome (Nordstrom and Dasgupta, 2006). The relative replication time of chromosomal loci can be determined experimentally by measuring the locus copy number in replicating cells relative to non-replicating cells; an approach termed marker frequency analysis. Recent technical advances have allowed genome-wide application of marker frequency analysis using microarrays or deep sequencing. Such experimental measures of genome replication can identify deviations from the predicted marker frequency that indicate perturbations to genome replication, such as DNA damage (Sangurdekar *et al*, 2010), chromosomal fragmentation (Kouzminova and Kuzminov, 2008), regions of variable fork velocity (DeMassy *et al*, 1987; Srivatsan *et al*, 2010), chromosomal rearrangements (Skovgaard *et al*, 2011), or pathological consequences of replication termination (Rudolph *et al*, 2013). However, in order to fully analyse the consequences of changes to replication dynamics in cells with distortions of DNA replication, we need a robust understanding of the parameters that define the experimentally obtained replication profiles.

**Figure 1.**
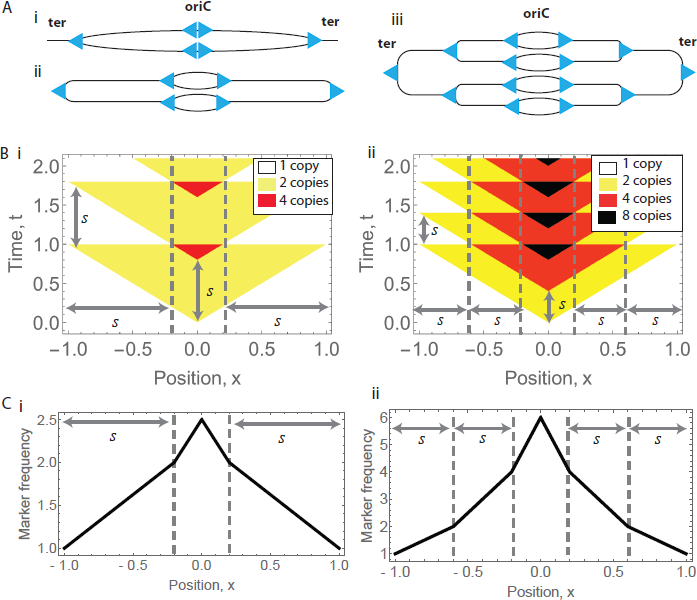
Representations of the replication program for *s* = 0.8 (left column) and *s* = 0.4 (right column). Position (*x*) unit is replichore length from origin to terminus. Time (*t*) and periodicity (*s*) units are time required to replicate the length *x* = 1 under assumption that fork velocity is constant. A Schematics of chromosome configuration at the initiation time (i) or termination time (ii and iii); the DNA is shown as black lines and the advancing replication forks are shown as triangles; *oriC* is position of the natural replication origin, *ter* indicates replication termination sites. Chromosome is linearised with respect to termination site. B Spatiotemporal representation as a ratio of copy number at *x* to copy number at *x* = *−*1.] C Model predicted marker frequency profiles.

A model for DNA replication during the bacterial division cycle was proposed by Cooper and Helmstetter (Cooper and Helmstetter, 1968). In this model, the average frequency of a particular locus per cell (e.g. a marker) can be predicted by taking into account the duration of chromosome replication, the time from the completion of chromosome duplication to complete cell division, the cell doubling time and distance between the loci and the origin. The model assumes that replication is initiated at every doubling in cell mass. A number of mathematical models have been formulated to describe other biological processes related to or influencing replication: simulation of gene expression in exponentially growing bacterial cells (Luo *et al*, 2013; Gomez *et al*, 2014), computation of theoretical DNA histograms for flow cytometry (Stokke *et al*, 2012), simulation of the bacterial cell division cycle (Zaritsky *et al*, 2011), the average cell oscillations in DnaA-ATP/DNA (Grant *et al*, 2011), or even a wide variety of 128 models that make di˙erent assumptions about the regulation of replication (Creutziger *et al*, 2012), but none of these studies considered the genome-wide parameters of replication.

In the present work, we have developed a new mathematical model for concurrent rounds of DNA replication. The model derivation is based on a spatiotemporal representation of the replication program (Retkute *et al*, 2012; Baker and Bechhoefer, 2014; Retkute, 2014). We test the proposed model on experimental data from the bacterium *E.coli* (Guzman *et al*, 2002; Riber *et al*, 2006; Skovgaard *et al*, 2011; Rudolph *et al*, 2013; Maduike *et al*, 2014). We found that a steeper gradient in replication profile is to be expected in the origin proximal region due to overlapping cell cycles. Furthermore, we extend this model for the example of DNA replication in *E.coli* with two origins (Wang *et al*, 2011; Rudolph *et al*, 2013) to gain valuable biological insight into this perturbed system. We found that fork velocity was reduced during times when there were twice as many forks as normal and synthesis was even slower for forks moving in a direction opposite to the normal direction of replication. As a consequence, a double origin strain grows slower compared to cells with a single origin. Finally, the ectopic origin may start ahead of the natural origin, but this has no e˙ect on the time required to complete synthesis.

## Results

### Modelling concurrent rounds of DNA replication

The initiation of DNA replication in fast growing bacteria occurs prior to the completion of the previous round of replication resulting in concurrent rounds of synthesis. Therefore, we define a parameter *s* as the periodicity of replication initiation, i.e. the time between two concurrent rounds of replication normalised by the time required to replicate the whole chromosome (Fig 1 and Supplementary Fig S1; for details see Materials and Methods). The model is defined in dimensionless variables: the chromosome positions are scaled relative to the distance between *oriC* and *ter*; while time, periodicity and di˙erence in initiations between origins, are scaled by the time required to replicate half of the chromosome. Assuming that the replication fork velocity is constant from initiation to termination, the periodicity can describe both temporal coordination of replication initiation and relative location of forks along the chromosome.

Figure 1 shows three representations of bacterial chromosome replication: schematics of chromosome replication configuration, spatiotemporal representation, and predicted marker frequency profiles; graphs are shown for two (*s* = 0.8, left panel) and three concurrent rounds of replication (*s* = 0.4, right panel). The marker frequency at a position *x* is the average number of copies in a population. The origin proximal regions have the highest copy number, and sites at *−*1 + *s* and 1 *− s* have exactly two copies, as these are the positions on the genome where forks from the subsequent round of replication are located when forks from the current round reach the *ter* site.

### Replication in wild-type *E. coli*

Our mathematical model predicts that replication profiles from fast growing bacteria will have at least one change in the slope, with the steepest replication profile gradient in the origin proximal regions. This is in contrast to other models which predict a continuous variation in slope approximating to an exponential decay (Cooper and Helmstetter, 1968). To test whether this model prediction is accurate, we fitted the model to 5 experimental data-sets (see Materials and Methods; Fig 2). Using only a single parameter (*s*), our model was able to give a fit to experimental data comparable or better than the Cooper and Helmstetter model (Cooper and Helmstetter, 1968), which requires knowledge of at least two cell cycle parameters (doubling time and time required to complete synthesis). Parameters for both models were fitted by minimising a mean squared error (Supplementary Table S1).

**Figure 2.**
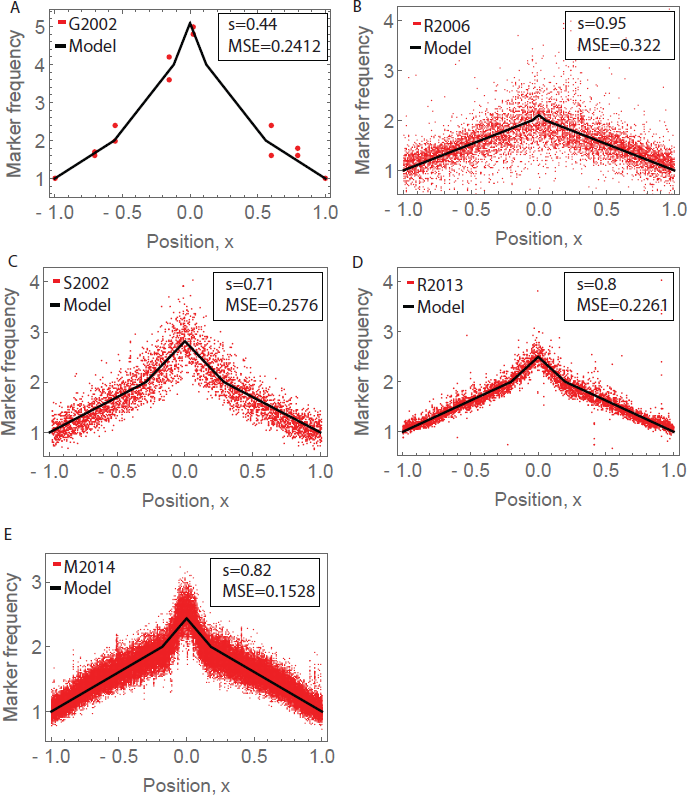
Model fitting for bacteria with one origin. Each panel shows fitted *s* value and mean squared error (MSE). Panels have di˙erent range of marker frequency with higher values corresponding to a smaller periodicity. Marker frequency profiles and model fits: A. Guzman *et al*, 2002, B. Riber *et al*, 2006, C. Skovgaard *et al*, 2011, D. Rudolph *et al*, 2013, E. Maduike *et al*, 2014.

### Replication in *E. coli*with two origins

Initiation of DNA synthesis occurring at multiple sites along a chromosome has been studied in *E.coli* by constructing a strain with two replication origins: the natural origin *oriC* and an ectopic origin *oriZ*, inserted 1 *Mb* away from the normal origin on the right replichore (Wang *et al*, 2011). Figure 3A shows the marker frequency profile (Rudolph *et al*, 2013) and the fitted model with assumption that both origins have identical properties. The fit to the experimental data is good, considering the simplicity of the model. However, the model shows some deviations from the marker frequency profile, in particular the peak heights and the slopes in the *oriC* and *oriZ* proximal region.

**Figure 3.**
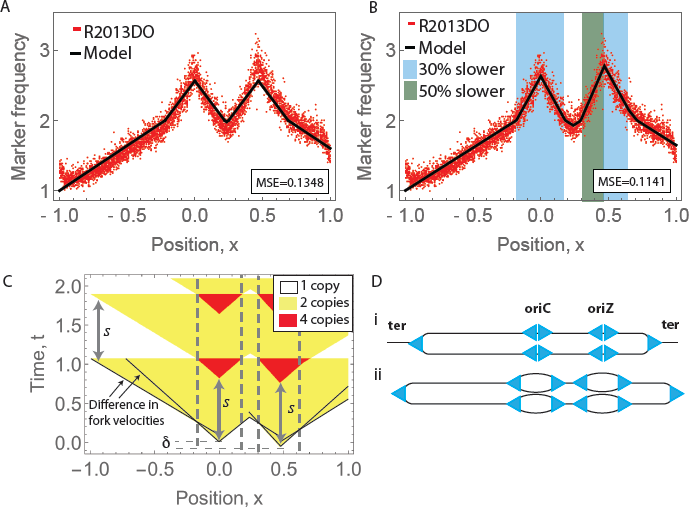
Model fitting for bacteria with two origins. A Marker frequency profile and model fit for *s* = 0.76 and no additional assumptions. B Marker frequency profile and model fit for *s* = 0.82, *δ* = 0.06 and reduced fork velocity (in blue and green regions). C Spatiotemporal representation of the replication program shown in B. Black lines indicate fork movement under various velocities. D Schematics of chromosome configuration at the initiation time (i) or termination time (ii and iii); the DNA is shown as black lines and the advancing replication forks are shown as triangles; *oriC* is position of the natural replication origin, *oriZ* is position of the ectopic replication origin„ *ter* indicates replication termination sites. Chromosome is linearised with respect to termination site. The maximum number of replication forks is 10.

In order to improve the fit, we refined our model to test our assumptions and gain biological insight into DNA replication in this two origin system. We allowed (i) forks to move with di˙erent velocities, and (ii) one of two origins to initiate earlier than the other. Our refined model gives a significantly better fit to the experimental data and provides evidence to support the following perturbations to DNA replication resulting from the addition of the ectopic origin (Fig 3B and C, Supplementary Fig S2). First, the model predicts that replication forks proximal to *oriC* and *oriZ* are slower than forks proximal to the termination site. A reduction in speed by 30% gave an significantly improved fit. The spatiotemporal representation indicates that fork velocity reduction occurs during times when there are twice as many forks compared to cells with a single origin (Fig 3C). Following the termination event between *oriZ* towards *oriC* the number of replication forks is comparable to that in wild-type cells and this is concomitant with a change to faster fork velocities. Second, the replication fork proceeding from *oriZ* towards *oriC* is further reduced in velocity (total 50% reduction relative to forks proximal to the termination site) and it is noteworthy that this fork is moving against the direction of highly transcribed genes. Third, one of the explanation for the di˙erence in peak heights in the *oriC* and *oriZ* proximal region (Supplementary Fig S3) may be that initiation at the ectopic origin occurs before initiation at the natural origin (population average *δ* = 0.06, i.e 6% of the time required to replicate the whole chromosome).

By directly comparing the experimentally measured profile with the profile based on the simple model, we were able to uncovered how cells respond when an extra replication origin is introduced to the chromosome. Most of the deviations can be readily explained by biological features which may be expected − activation of the ectopic origin will require additional replication forks, and some of these forks will move against the direction of highly transcribed genes. Our accurate mathematical model should be useful tool in experiments where perturbations to replication process are not known in advance, for example, in the presence of defects in nucleic acid metabolism.

## Discussion

We have formulated a mathematical model that allows interpretation of DNA replication initiation dynamics from marker frequency profiles. We have successfully applied this model to study replication dynamics in wild type *E. coli* cells. Analysis of this mathematical model shows that the replication profile is composed of a set of varying slopes (replication profile gradients). This has an important biological implication − the change in the gradient in the *oriC* proximal region is predicted as a consequence of overlapping cell cycles, not as a consequence of variability in replication fork velocity. The increased steepness of the slope of the profile 0.3 *Mb* on each side of *oriC* previously has been attributed to either replication forks moving slower over this interval, or some unusual perturbation of replication initiation (Maduike *et al*, 2014) (Fig 2E). This region coincides with a marker frequency gradient change predicted by the model without the need to invoke a variable fork velocity. The model precisely predicts the coordinates where the slope of replication profile changes: it is (1 *−* 0.71) *×* 4.63 *Mb/*2 = 0.42 *Mb*. Therefore, our mathematical model is able to explain that a steeper gradient is to be expected due to the overlapping cell cycles without assuming any perturbation of the speed of synthesis. The model can describe replication in other bacteria such as *Bacillus subtilis*, as shown in Supplementary Fig S4 (Kono *et al*, 2014).

To maintain constant genomic content at division, the doubling time should be directly proportional to the periodicity of initiation. If the time required to replicate the chromosome, *C*, is known, then doubling time *τ* = *s×C* (where *τ* and *C* are in minutes and *s* is unitless). This means that the periodicity of initiation is what sets the cell division time. The growth rate (doublings per minute) can be calculated as 1*/*(*s × C*).

We assumed that there is little variation in the periodicity of origin firing. This implied a discrete location for the change in gradient in replication profiles (Fig 2). If this assumption is relaxed and initiation can occur in a window [*s −* Δ*s, s* + Δ*s*], replication profiles will have a smoother transition between gradients (Supplementary Fig S5). However, allowing less defined periodicity does not improve the model fit (Supplementary Table S2). This suggests that there is a strict periodicity in replication initiation, which is consistent with known feedback loop that regulates initiation time (Browning *et al*, 2004).

The comparative analysis of our simple model with the data obtained in a double *ori* background reveals di˙erences, which allows us to gain insights into the processes that govern DNA replication when replication is perturbed. Any deviation from the model can indicate alterations to replication parameters. By adjusting the model assumptions we can add extra parameters, analyse their influence and uncovered how cells respond to the presence of two origins. The model deviations predict variability of the following parameters: reduced fork velocity, longer chromosome synthesis time, and possible earlier initiation of the ectopic origin.

The overall reduction in fork velocity proximal to *oriC* and *oriZ* could be a consequence of limited dNTP concentrations if there are insu˚cient compensatory mechanisms to supply the increased number of forks resulting from the addition of the ectopic origin (Nordman and Wright, 2008). For example, in cells with a single origin undergoing concurrent rounds of replication the total number of replisomes per cell is limited to approximately 10 (Kelman and O’Donnell, 1995). The maximum number of replisomes per genome is 10 in *E. coli* with two origins (Figure 3D). Therefore, the rapid speed of DNA synthesis may be compromised due to the intracellular scarcity of dNTP molecules.

The further reduction in fork velocity upstream of *oriZ* is likely to be due to head-on collisions between this replication fork and transcription complexes. There are 36 mRNA promoters with transcriptional start sites encoded on the leading strand in this region (Supplementary Information, Hershberg *et al*, 2001). The replisome may stall when it encounters a RNA polymerase moving in the opposite direction but eventually resumes elongation after displacing the RNAP from DNA (McGlynn *et al*, 2012). Similar behaviour was observed when transcription was reversed by inverting a quarter of the *B. subtilis* chromosome downstream of the origin, this led to 30% decrease of replication rate (Srivatsan *et al*, 2010). This implies that the site of replication termination between *oriC* and *oriZ* is dictated not solely by replication timing and fork velocity, but is modulated by transcription (Rudolph *et al*, 2013).

We found a slower cell doubling for the double origin strain compared to wild-type cells: 23 *min* in rapidly growing bacteria with one origin, and 25 *min* in rapidly growing bacteria with two origins (Supplementary Information). This is in agreement with experimentally measured doubling times in rich media, which were 23 *min* for the single origin strain and 24 *min* for the two origin strain, however, the di˙erence observed experimentally is not statistically significant (Wang *et al*, 2011).

Fitting the modelled replication profile to the experimental data indicated that initiation at the ectopic origin may occur before initiation at the natural origin, which could be a consequence of the di˙erences in accessibility for DnaA binding due to nuclear di˙usion (Berg, 1984) or macromolecular crowding (Akabayov *et al*, 2013). However, this does not a˙ect the time required to complete synthesis, as the distance travelled by the rightward moving fork initiated at *oriZ* is about 50% shorter then the distance traveled by the leftward moving fork initiated at *oriC* (1.25 *Mb* and 2.31 *Mb*). So the former reaches *ter* site earlier, but is blocked by terminator proteins and has to wait in the *ter* region. Under normal conditions, the left and right replicons are approximately equal in size, so the di˙erence in initiation timing may indicate there is a mechanism which tries to regulate time spend by replisome in the termination zone. Further experimental data is required to verify if this e˙ect is a consequence of the particular growth conditions or bacterial strain.

We expect that our proposed mathematical formulation can be adopted for various DNA replication alterations. Although the model is formulated for a bacterial genome, it is relevant to replication in archaea (Hawkins *et al*, 2013), the development of genetic instability in tumour cells due to rereplicating DNA segments (Vaziri *et al*, 2003), plant cell growth during endoreplication cell cycle (Edgar and Orr-Weaver, 2001) or replication in polytene nurse cell from *Drosophila* (Hammond and Laird, 1985). By focussing on specific and more specialised aspects of chromosome replication, our model can be further refined to gain deeper biological insight into the regulation of DNA replication. With deep sequencing technologies providing cheaper and higher-resolution profiles of whole genome replication, there is an increasing requirement for mathematical models to maximise our biological understanding of these fundemental cellular processes.

## Materials and Methods

### Marker frequency

We have analysed five experimental data-sets for *E. coli* with one origin, and one experimental data-set for *E. coli* with two origins. Details on the data, method and growth conditions are given in Table 1. The chromosome positions were scaled relative the distance between *oriC* and *ter* (Fig 2).

**Table 1:**
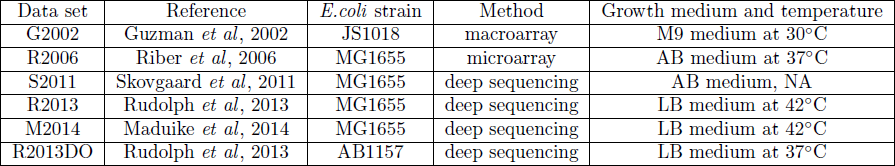
Summary of *E. coli* data-sets used for the analysis.

| Data set | Reference | *E.coli* strain | Method | Growth medium and temperature |
| --- | --- | --- | --- | --- |
| G2002 | Guzman *et al*, 2002 | JS1018 | macroarray | M9 medium at 30°C |
| R2006 | Riber *et al*, 2006 | MG1655 | microarray | AB medium at 37°C |
| S2011 | Skovgaard *et al*, 2011 | MG1655 | deep sequencing | AB medium, NA |
| R2013 | Rudolph *et al*, 2013 | MG1655 | deep sequencing | LB medium at 42°C |
| M2014 | Maduike *et al*, 2014 | MG1655 | deep sequencing | LB medium at 42°C |
| R2013DO | Rudolph *et al*, 2013 | AB1157 | deep sequencing | LB medium at 37°C |

#### Mathematical model for DNA replication in bacteria with one origin

Mathematical model for DNA replication during overlapping cell cycles in bacteria is based on the spatiotemporal representation of the replication program (Retkute *et al*, 2012; Baker and Bechhoefer, 2014; Retkute, 2014). Replication initiates at replication origin *oriC* and proceeds bidirectionally around the chromosome to a site opposite *oriC* called the replication terminus *ter* (Fig. 1A). For simplicity we virtually linearise the chromosome by putting replication origin at position *x* = 0, and normalise its length so that the distance between *oriC* and *ter* is one arbitrary unit (a.u.). The chromosome is divided into a left replichore at *x* ∊ [*−*1, 0] and right replichore at *x* ∊ [0, 1]. Each replication initiation event creates two replication forks which move with a constant velocity *v* to the opposite directions away from *oriC*. Replication timing *R*(*x*) is defined as the time at which the locus *x* is replicated. For a single initiation event the replication timing at the site *x* is given by (de Moura *et al*, 2010; Yang *et al*, 2010):

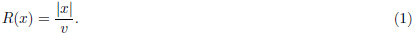

If we define a time unit one as the time required to complete synthesis of the whole chromosome, and assume that the replication fork velocity is constant from initiation at *oriC* to termination at *ter*, we obtain from Eq.(1) that 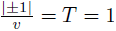,therefore *v* = 1.

We defined the periodicity of initiation, *s*, as a time interval at which new initiation events occur. Therefore replication timing of *k^th^* initiation is *R_k_*(*x*) = *|x|* + (*k −* 1)*s*. When a new round of replication is initiated, forks from the previous round have moved *v × s* = *s* away from *oriC*, i.e. the distance is exactly *s*.

Based on the spatiotemporal representation of the replication program we can derive an analytical expression for marker frequency at a position *x* which depend only on the parameter *s*. Marker frequency profile for replication case where up to two replication rounds occur is given by the following equation:

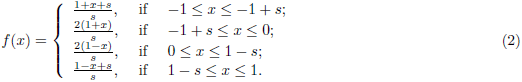

Derivation of the model is given in Supplementary Information.

### Mathematical model for DNA replication in bacteria with two origin

If there is more then one origin present on the chromosome, then the replication time of a given position in the chromosome is found by evaluating

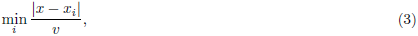

where *x_i_* is the positions of the origin *i* (de Moura *et al*, 2010; Yang *et al*, 2010). The marker frequency is calculated from the spatiotemporal representation of the replication program (details in Supplementary Information).

### Parameter estimation

Parameters were fitted by minimising a mean squared error between model predicted marker frequency, *f_i_*,and experimentally measured marker frequency, *d_i_*:

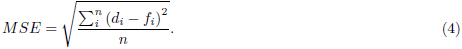

## Acknowledgements

The authors thank Prof Anders Lobner-Olesen, Prof Ole Skovgaard and Dr Kono Nobuaki for providing experimental data.

## Author Contributions

R.R., C.A.N, C.R. and M.H. designed the study. R.R. developed the mathematical model and conducted calculations. R.R., C.A.N, C.R. and M.H. wrote and edited the manuscript.

## Supplementary Information

### Modelling replication in bacteria with one origin

Our model has the following assumptions: (i) chromosome is linearised at *ter* site; (ii) *oriC* is positioned at *x* = 0; (ii) chromosome length is normalised in such a way that the distances between *oriC* and *ter* on left and right replisomes are equal to 1; (iii) replication forks move with a constant speed; (iv) time required for replication forks to travel from *oriC* to *ter* is *T*; so that the time unit is time required for replication forks to travel from *oriC* to *ter* with no perturbations of replication; (vi) the time interval at which new initiation events occur is *s* (the periodicity of initiation). The overlapping cell cycles will satisfy condition: *s < T*.

We define the rule of replicating position *x* as *R*(*x*). This rule for assumptions (i)-(iii) becomes *R*(*x*) = *|x|*. We start from a single copy of the genome, and at times *t_i_* = *is*, *i* = 1, 2*, …*, a new replication round is initiated at *oriC*. The number of replication initiations occurring before *t* can be found as:

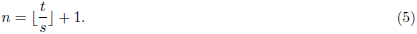

Time when a genomic site *x* is replicated by forks which have initiated at round *i^th^* is given by *R*(*x*)+(*i−*1)*s* and at this time 2*^i− 1^* new copies are made.

Then the copy number at position *x* at the time *t* is equal to:

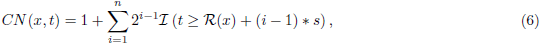

where *I*(*x*) = 1 if condition *x* is satisfied and zero otherwise.

Figure S1A shows the dynamics of copy number at positions *x* = 0 and Figure S1B at *x* = *−*1. It can be seen that even after a few rounds of replication, the copy number increases exponentially.

**Figure S1.**
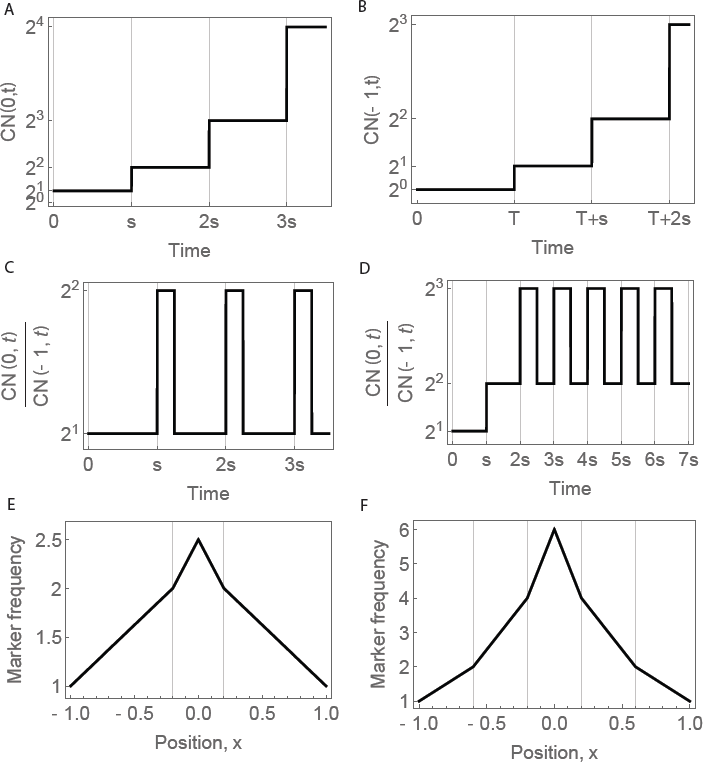
Model is formulated based on periodicity *s*. A The dynamics of copy number at position *x* = 0. B The dynamics of copy number at position *x* = *−*1. C Ratio of copy number at *x* = 0 to copy number at *x* = *−*1 as a function of time for *s* = 0.8. D Ratio of copy number at *x* = 0 to copy number at *x* = *−*1 as a function of time for *s* = 0.4. E Model predicted replication profiles for *s* = 0.8. F Model predicted replication profiles for *s* = 0.4.

What we actually are interested is not the value of *CN*(*x, t*) itself, but its value with respect to a reference position along a genome, i.e.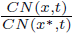. If we follow the changes in this ratio over the time, we can see that this is a periodic function with period *s*. Figure S1C shows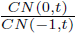as a function of time for *s* = 0.8 and Figure S1D for *s* = 0.4. This function becomes periodic for *t*≥ *T*.

Therefore, to calculate marker frequency, we need to find the average copy number over a single period. We choose this interval to be *t* ∊ [*T, T* +*s*], i.e. the time forks from the first round of replication have reached *ter* sites. Then marker frequency can be calculated using the following expression:

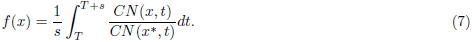

For further analysis we assume that 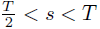(i.e. there are up to two rounds of replication). We choose *ter* as the reference point, i.e. *x^*^* = *−*1. It can be seen from Fig. 1A that for *t* ∊ [*T, T* + *s*], *CN*(*x^*^, t*) = 2. Next, for each position *x* we find the largest integer *K* such that satisfies *T < R*(*x*) + *sK ≤ T* + *s*. Then, we can rewrite Eq.(6) as:

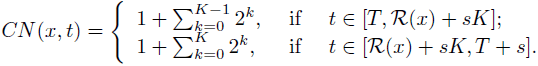

Now we can integrate Eq.(7):

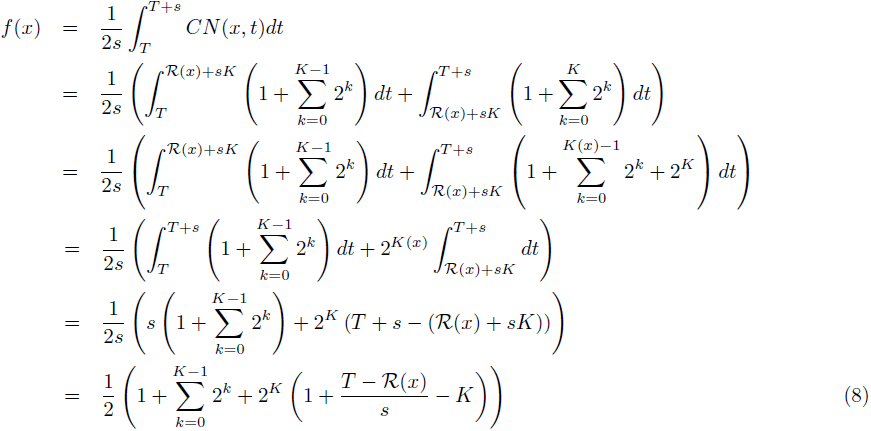

We can derive an analytical expression for marker frequency by taking into account that *T* = 1, *R*(*x*) = *|x|* and *K* either equal to one or two:

1. For *−*1 *x −*1 + *s*, we have that *K* = 1 and Eq.(8) gives:

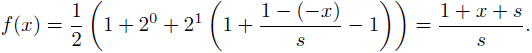
2. For 1 *− s x* 1, we have that *K* = 1 and Eq.(8) gives:

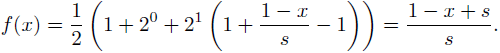
3. For *−*1 + *s x* 0, we have that *K* = 2 and Eq.(8) gives:

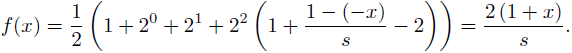
4. For 0≤ *x*≤ 1 *− s*, we have that *K* = 2 and Eq.(8) gives:

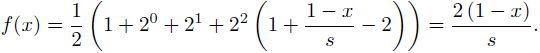

Therefore, marker frequency profile for replication case where up to two replication rounds occur is given by the following expression:

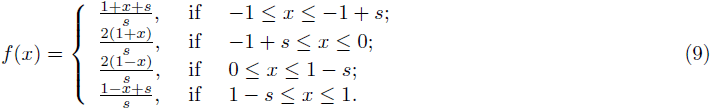

### Model with less defined periodicity

We assume that initiation can occur within intervals [*i × s* − Δ*s, i × s* + Δ*s*], *i* = 0, 1*, …*. Then copy number is defined by the following mathematical expression:

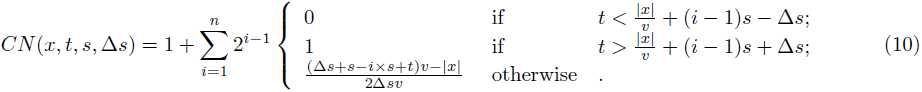

where *I*(*x*) = 1 if condition *x* is satisfied and zero otherwise. Supplementary Fig. S4 shows copy number, ratio between copy number at *x* = 0 and *x* = *−*1, and replication profiles for *Δs* = 0.05 and *Δs* = 0.1 and two values of *s* = 0.8 and *s* = 0.4.

**Figure S3.**
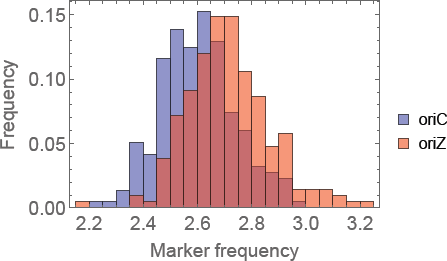
Distribution of marker frequency in the *oriC* and *oriZ* proximal region (*±*0.05 from the origin position). Mann-Whitney U test indicates statistically significant di˙erences between the two peak heights (*p* = 9.9351 *×* 10*^−14^*).

**Figure S4.**
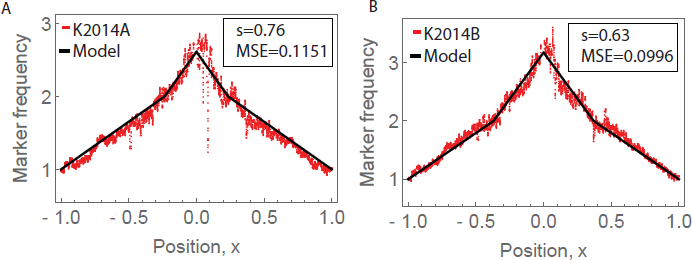
Model fitting for *Bacillus subtilis* (Kono *et al*, 2014). Each panel shows fitted *s* value and mean squared error (MSE). Panels have di˙erent range of marker frequency with higher values corresponding to a smaller periodicity. Marker frequency profiles and model fits: A. K2014A: wild type strain, B. K2014B: *rtp*^+^ mutant.

### Modelling replication in bacteria with two origins

Any alterations to the replication program can be incorporated into the mathematical model by adjusting expression of *R*(*x*) and *T*, and then solving Eqs.(5)-(7).

If forks are moving with speed *w* in the vicinity of replication origin, and with a normal speed *v* = 1 elsewhere (i.e. *w≤ v*), then

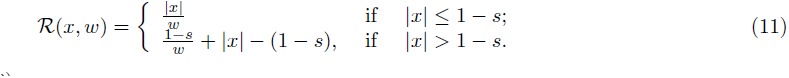

We can rewrite Eq.(6) as:

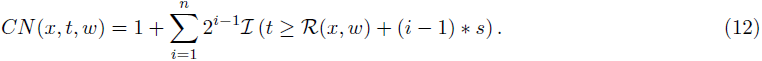

Now time required for the fork move from *oriC* to *ter* on the left replichore is 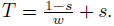.

The marker frequency profile for two origins with similar properties and no alteration in replication program can be calculated by evaluating the integral Eq.(7) with the following function:

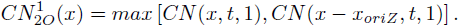

Spatiotemporal representation and marker frequency profile together with the experimental data R2014DO for the above case is shown in Fig. S2A. The model shows some deviations from the experimental data, in particular the peak heights and the slopes in the *oriC* and *oriZ* proximal region. To account for these observed di˙erences, we assumed that both pairs of forks move slower around replication origins. Then marker frequency profile is calculated as:

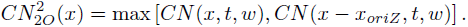

**Figure S2.**
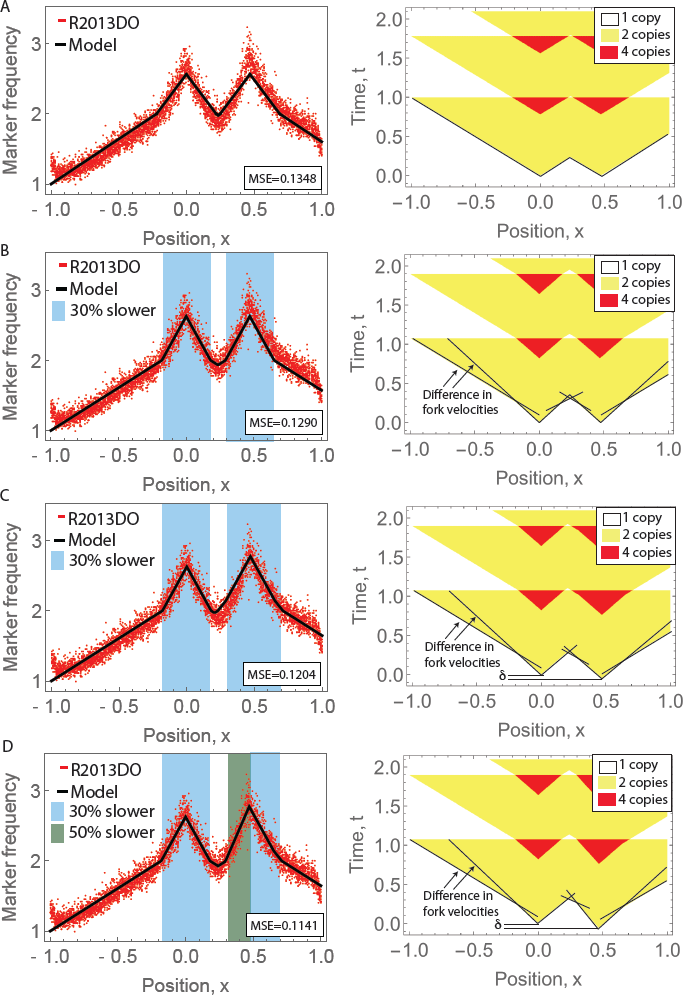
Model fitting for bacteria with one origin (left column) and spatiotemporal representation of replication program (right column). Black lines indicate fork movement under various velocities. A Model fit with no additional assumptions. B Model fit under assumption that forks are moving 30% slower. C Model fit under assumption that forks are moving 30% slower and *oriZ* initiates *τ* earlier in comparison with *oriC*. D Model fit under assumption that forks are moving 30% or 50% slower and *oriZ* initiates *τ* earlier in comparison with *oriC*.

**Figure S5.**
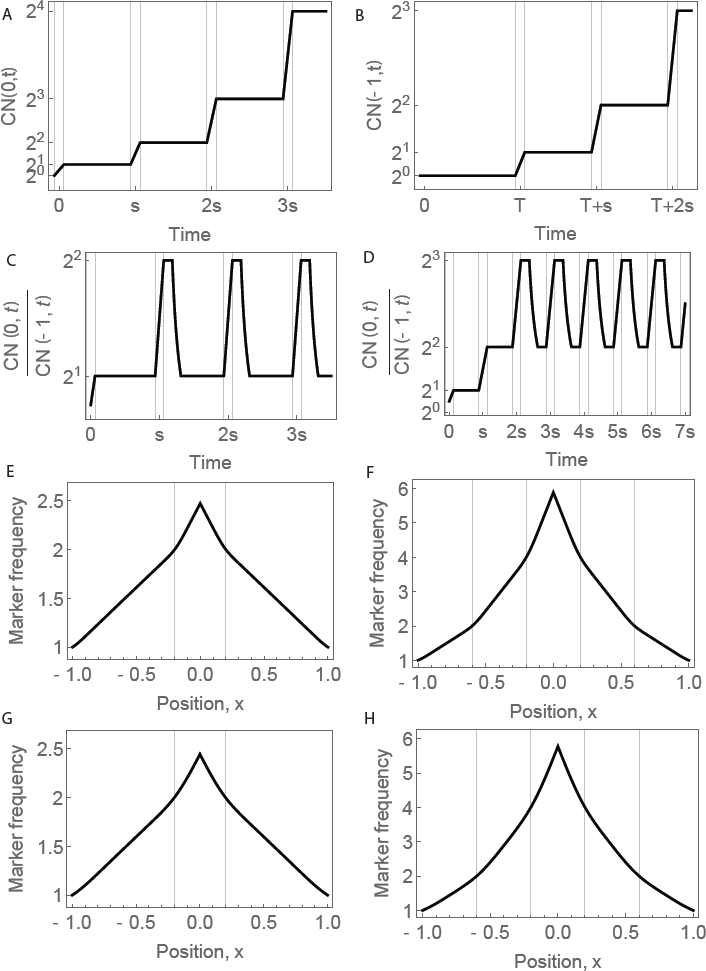
Model with less defined periodicity. In A-D horizontal lines show *i × Δs ± s*. In E-H horizontal lines show *−*1 + *i × s* and 1 *− i × s*. A The dynamics of copy number for Δ*s* = 0.05 at position *x* = 0. B The dynamics of copy number for Δ*s* = 0.05 *x* = *−*1. C Ratio of copy number for Δ*s* = 0.05 at *x* = 0 to copy number at *x* = *−*1 as a function of time for *s* = 0.8. D Ratio of copy number for Δ*s* = 0.05 at *x* = 0 to copy number at *x* = *−*1 as a function of time for *s* = 0.4. E Model predicted replication profiles for Δ*s* = 0.05 and *s* = 0.8. F Model predicted replication profiles for Δ*s* = 0.05 and *s* = 0.4. G Model predicted replication profiles for Δ*s* = 0.1 and *s* = 0.8. H Model predicted replication profiles for Δ*s* = 0.1 and *s* = 0.4

Spatiotemporal representation and marker frequency profile for the above case is shown in Fig. S2B. Slower moving forks improved the model fit around *oriC* and on the whole left replichore, but yet the model fit is below data points in *oriZ* proximal region. Therefore, we assume that *oriZ* is activating *˝* time units earlier then *oriC*:

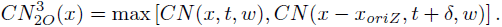

Spatiotemporal representation and marker frequency profile for the above case is shown in Fig. S2C. This has improved fit on the right of the *oriZ*, but the sloper on the left became too high. This means that forks are moving even slower in the region. So finally, to allow the heterogeneity in replication around *oriZ*, on the left forks move with velocity *u* and on the right with velocity *w*:

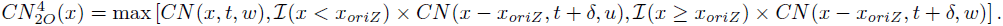

Spatiotemporal representation and marker frequency profile for the above case is shown in Fig. S2D.

### The Cooper and Helmstetter model

The bacterial division cycle can be partitioned into three phases: time before chromosome replication, *B*, duration of chromosome replication, *C*, and time from completion of chromosome duplication to complete cell division, *D*. The model of DNA replication during bacterial division cycle has been derived by Cooper and Helmstetter (Cooper and Helmstetter (1968)). Parameterization of the Cooper and Helmstetter (CH) model requires knowledge of the length of division cycle phases and the length of doubling time, *τ*. Marker frequency of a genome site at a position along chromosome, *x*, is given by:

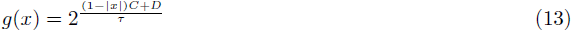

Supplementary Table S1 gives MSE when parameters were fitted to our mathematical model and CH model. For CH model we have minimised MSE with respect to a doubling time *˝*. As there is no division, we have set *D* = 0 min. For all data except Skovgaard *et al.* our proposed model outperformed CH model.

### mRNA promoters in the region of slower forks

Promoter (position on *E.coli* genome bp): *dnaKP1* (12047), *dnaKP2* (12123), *dapB* (28343), *carABP1* (29551), *carABP2* (29619), *carB* (30775), *caiF* (34218), *folA* (49799), *araC* (70223), *ilvIHP4* (85394), *ilvIHP3* (85420), *ilvIHP2* (85534), *ilvIHP1* (85597), *ftsQP2* (102742), *ftsQP1* (102867), *ftsZP4* (104636), *ftsZP3* (104693), *ftsZP2* (105045), *pdhR* (122034), *aceE* (122970), *lpd* (127720), *acnB* (131519), *htrA* (180844), *accA* (208413), *dnaQP1* (235934), *dnaQP2* (236017), *betT* (328645), *codB* (354108), *cynT* (357997), *fimBP2* (4538235), *fimBP1* (4538377), *deoP1* (4614248), *deoP2* (4614847), *deoP3* (4617136), *serB* (4622432), *trpR* (4630273).

### Cell doubling times

The replication initiation events occurred at similar intervals in *E.coli* with one origin (*s*_1_ = 0.82) and two origins (*s*_2_ = 0.80). However, our analysis indicate that the double origin strain grows more slowly: the counter-clockwise replichore has the same length as its counterpart in single origin strains (*L* = 2.32 *Mb*), but fork velocity was reduced in *oriC* proximal regions, which will increase the time required to replicate the replichore. If we assume that a nominal fork velocity is *v* = 0.081 *Mb/min*, the time required to replicate half of the chromosome is *C*_1_ = *L/v* = 2.32 *Mb/*(0.081 *Mb/min*) *≈* 28 *min*, while doubling time *τ*_1_ = *s*_1_ *× C*_1_ = 0.82 *×* 28 *min ≈* 23 *min*. In the double origin strain, the region *s*_2_ *× L* in counter-clockwise replichore is replicated at the nominal fork velocity *v*, while the rest of replichore is replicated by forks travelling 30% slower, i.e. with velocity 0.7*×v* (Fig. 3B). Therefore, the time required to replicate the double origin strain is *C*_2_ = *s*_2_ *×L/v* +(1*−s*_2_)*×L/*(0.7*v*) = 0.8*×*2.32 *Mb/*(0.081 *Mb/min*)+0.2*×*2.32 *Mb/*(0.7*×*0.081 *Mb/min*) = 31 *min*, while doubling time is *τ*_2_ = *s*_2_ *× C*_2_ = 0.8 *×* 31 *min ≈* 25 *min*.

**Supplementary Table S1** Fitting model for bacteria with one origin: MSE comparison between our proposed model and The Cooper and Helmstetter model.

**Supplementary Table S2** Fitting model with less defined periodicity for dataset R2013 (Rudolph *et al*, 2013 and dataset M2014 (Maduike *et al*, 2014).

